# Characterization of sub-tropical maize (*Zea Mays* L.) inbred lines for the variation in kernel row numbers (KRN)

**DOI:** 10.1101/2022.09.23.509129

**Authors:** Ganapati Mukri, RN Gadag, Jayant S Bhat, T. Nepolean, Navin C Gupta, Shikha Mittal, ML Nithyashree, Ramesh Kumar, Digvender Pal

## Abstract

Genetic dissection of high KRN trait and cob length has been undertaken by several researchers resulting in identification of loci controlling the traits. Further fine mapping of QTLs controlling KRN and cob length and TILLING strategies identified the underlying genes. All these studies are used temperate maize genotypes which are of limited use to researcher in sub-tropical region, and the sub-tropical maize production systems. Present investigation explores the availability of genetic regions responsible for KRN in sub-tropical maize germplasm. A total of 280 subtropical maize germplasm was analyzed for their KRN variation and selected 45 stable lines were subjected to molecular characterization using genes linked to KRN in maize. Diversity analysis was also performed to understand the possible association of character with the KRN gene in deciding its variation in the given population. It was revealed that four genes showed highest probability of influencing KRN traits in these tropical maize inbred lines. The remaining genes not establishing specific pattern of association with high and low KRN genotypes may need further study on its allelic variation.

## Introduction

Maize is known as a queen of cereal by virtue of various factors and features like versatility, amenability for diverse uses, greater adaptability, its C4 nature and high yield potential. In India, maize is the third most important food crops after rice and wheat and produces 27.82 million tonnes (MT) maize from the crop cultivated in an area of 9.2 million hectares (mha) with a productivity of 3 tonnes/ha (FAOSTAT, 2018). India contributes 2-3% of the total maize production worldwide and amongst the top five maize exporters of the world, contributing almost 14% of the total maize exported to different countries around the globe. The demand for maize as food, derived food, feed and fodder is increasing in tune with expanding global population (Dass et al. 2009). Maize plays an important role in maintaining food security, prompts development of animal husbandry and maize based industries (Mukri et al. 2018). Thus, consistently improvement of grain yield is an important objective for maize breeders and also a challenge to increase the productivity of maize (Yi Q et al. 2019). To meet these demand prospective and potential requirements, breeder and geneticists working in maize have huge challenge of enhancing the productivity vis-à-vis with the decreasing natural resources. Understanding the genetic basis of grain yield is necessary to guide breeding efforts towards the development of high-yielding hybrids along with their specific adaptations (Rakshit et al. 2012). Maize yield a complex trait and it is the cumulative effect of expression of its component traits such as cob length, cob girth, kernel row number, kernel per row and shelling percentage and their interaction with environments (Bommert et al. 2013, Calderón et al. 2016). As one of the major mutations, increased kernel row number, which underwent a dramatic change from two rows in teosinte to more than eight rows in modern maize, is directly associated with the increased grain yield (Clifford F. Weil & Rita-Ann Monde, 2007). Kernel row number (KRN) is one of the most important yield contributing traits and a major breeding target to increase maize productivity (Liu L, et al, 2015). The variation in KRN of inbred lines plays a major role in determining the *perse* plant yield of that hybrid for which they are contributing as parental lines. Over recent decades, the scientific community has been working to uncover the genetic architecture of grain yield and yield-related traits in maize *(Zea mays* L.) (Shen et. al, 2019). Better understanding of the genetic architecture of KRN is required to establish breeding programs aiming at improving *per se* yield of maize hybrids. However, current knowledge about the genetic control of the maize KRN was obtained mainly from genetic assays of inflorescence mutants. For example, *thick tassel dwarf 1* (*td1*) (Bommert et al. 2005) and *fasciated ear 2* (*fea2*) (Taguchi-Shiobara et al. 2001; Bommert et al. 2013) are involved in meristem maintenance. The *ramosa* mutants can increase the indeterminacy of lateral organs, which transforms the determinate spikeletpair meristems into branches (Bortiri et al. 2006; Gallavotti et al. 2010; Satoh-Nagasawa et al. 2006; Sigmon and Vollbrecht 2010). Similarly, *Suppressor of sessile spikelets 1 (Sos1)* controls meristem determinacy to produce single instead of paired spikelets in the inflorescence, thereby decreasing the KRN in the ear (Wu et al. 2009). The dominant *Corngrass1* (*Cg1*) mutant encodes two tandem *zma-miR156* genes and leads to a small ear lacking an ordered kernel row and unbranched tassel (Chuck et al. 2007). It is pertinent to note that Indian maize germplasms are not subjected to comprehensive and systematic studies in respect of KRN traits. Generating such information about KRN, could be better exploited in the crop improvement program in general and hybrid development in particular. Therefore, this study is an effort to characterize Indian maize inbred lines, which would be helpful in further dissecting the genetics of KRN traits.

## Materials and methods

### Maize germplasm

A set of 280 sub-tropical maize inbred lines were evaluated consecutive two seasons, during *kharif* 2017 and *rabi* 2017-18, at ICAR-Indian Agricultural Research Institute, New Delhi, under augmented design. The cob parameters such as cob length, cob girth, kernel row number, kernels/row and shelling percentage were recorded and analyzed. The population showed variability for the studied traits (Supplementary Table 1). The cob length ranged from 3.2-20.80 cm in *kharif* and 6.75-19.60 cm in rabi and kernel row number, which ranged from 10-26, in both the studied seasons, were considered as basic criteria to narrow-down genotypes to total 80 for further characterization. During *kharif* 2018, these selected 80 inbred lines were evaluated in augmented design and were characterized for kernel and cob parameters. The mean and range for the traits indicated that the genotypes have significant variability for kernel row number and cob length (Supplementary Table 2). The phenotypic coefficient of variation and genotypic coefficient of variation showed that these traits were less influenced by environmental factors. Among the 80 genotypes, based on the KRN, total 16, 42 and 22 genotypes were grouped as low (<=14 KRN), medium (16-18 KRN) and high (>18 KRN) KRN genotypes respectively. Finally, 15 genotypes each from low, medium and high category were characterized comprehensively for kernel row number and associated traits across the three different environments *viz*., Location 1. *Kharif* season, 2018 at IARI, New Delhi. Location 2. *Rabi* season, 2018-19 at IARI New Delhi. Location 3. *Rabi* season, 2018-19 at RRC-Dharwad.

### Selection of genes for KRN variability

Out of several genes known for having role in influencing kernel row number, there is a need to choose specific genes which could be informative. A total of 13 genes controlling KRN traits, namely., *fea2* (Taguchi-shibara et al, 2001), *ct2* (Bommert et al, 2013), *td1* (Bommert et al, 2005), *Ids1*(Chuk et al, 1998), *Kn1*(Brown et al, 2011), *zfl1* and *zfl2* (Bomblies et al, 2005), *Fea3* (Byoung et al, 2016), *fea4* (Pautler et al, 2015), *cg1*(Chuck et al, 2007), *tsh4* (Chuck et al, 2010), *ub2* and *ub3*(Chuck et al, 2014) were selected (Table.1) for analyzing their sequence variation and possible association with the KRN across the 45 selected genotypes. The gene sequences were obtained from the public domain; https://www.maizegdb.org/.

**Table:1.**
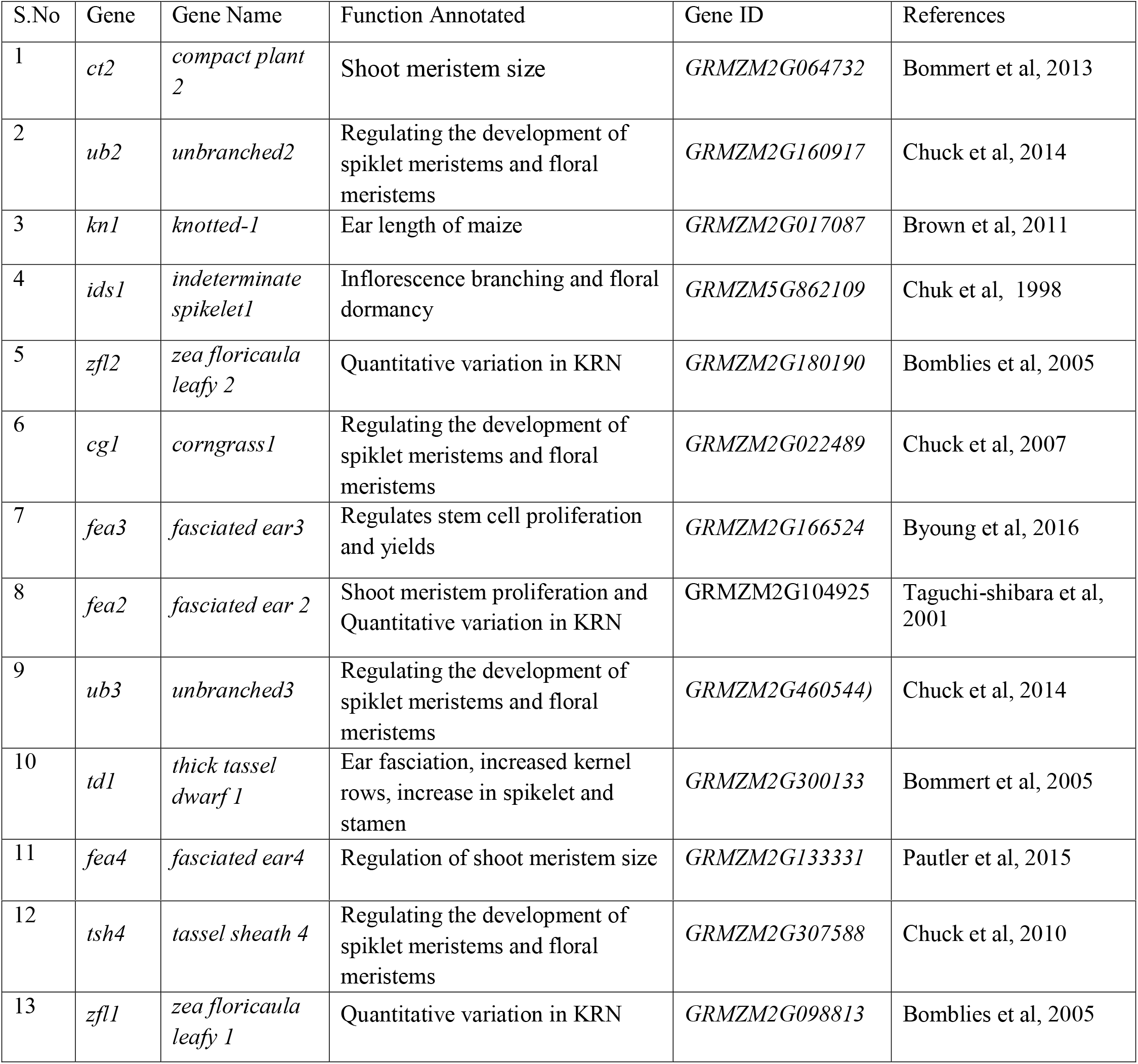
Details of the gene used for characterizing the inbred lines

### Isolation of genomic DNA and purification

The genomic DNA was isolated manually from the young leaves of the individual plants by following the CTAB method as described by Saghai-Maroof et. al. (1984). Isolated crude genomic DNA was purified by PCI (phenol-chloroform-isoamyl; 25:24:1) method and was diluted to the concentration of 500ng/μl after checking their quality by electrophoresis on 0.8% agarose gel.

### Primer designing and PCR amplification

The primers were designed manually from the flanking region of the gene sequence and evaluated them through DNA Club software (https://molbiol-tools.ca/molecular_biology_freeware.htm) with standard parameters. The gene length varied from 0.5-8.0 kb, hence, normal as well as long PCR amplification were done for the respective genes across the genotypes under investigation. The DNA polymerase ‘Takara PrimeSTAR GXL Taq’ was used for standard PCR to amplify the genes up to 3 kb in size whereas ‘NEB Long Amp’ was used to amplify the genes with more than 3 kb in size.

The PCR mixture for Takara PrimeSTAR GXL Taqbased gene amplification was constituted of 5X PrimeSTAR GXL Buffer (5 μ l), dNTP Mixture (2.5 mM each) (2 μl), primer 1 (10-15 pmol), primer 2 (10-15 pmol), Template (50ng), DNA Polymerase (0.5 μl) and Sterile distilled water (volume to 25 μl) with the thermal program of denaturation at 98°Cfor 10 second, annealing at 62°C for 15 seconds and extension at 68°C for 2 min.(min/kb) and the cycle was repeated in thermal cycler for 30. For NEB Long PCR, amplification was carried out using the PCR mixture composed of 5X Long Amp Taq Buffer (5μl), dNTPs (0.75 μl), Primer 1 (1 μl), Primer 2 (1 μl), NEB Long Amp Taq DNA Polymerase (2.5 U), Template 50ng and water (Volume up to 25 μl). Thermal program of the long PCR amplification was followed as per the manufacturer protocol.

### Statistical analysis

The analysis of variance (ANOVA) for additive main effect and multiplicative interaction (AMMI) and GGE biplot analysis were carried out using software Genstat. Diversity analysis by hierarchical clustering with Ward’s method was performed to understand the relatedness of the studied genotypes using SPSS software. Descriptive statistics and analysis of variance (ANOVA) were estimated using the software SAS 9.3 v (http://stat.iasri.res.in/). Logistic regression analysis was used to assess the relationship between the number of kernel rows as a binary variable (<=14 KRN=0 and >14 KRN=1) and KRN associated gene as explanatory variables. The following formula was used to estimate the probability of gene under investigation influences the kernel row number in maize inbred line by the maximum likelihood method (Hosmer and Lemeshow, 1989).

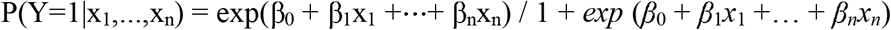

## Results

### Phenotypic variability, stability and diversity in selected maize germplasm

The ANOVA of AMMI analysis indicated that genotypes evaluated across the environments were significant among each other. The cob length, cob girth, kernel row number and grain yield showed significant G×E interaction (Table 2). Environment in which the genotypes were evaluated also were significant. The GGE biplot analysis indicated that three each genotype, AI 544, AI 521, AI 533 and AI 531, AI 545, AI 504 were specifically adapted to location 2 and location 3 respectively. The genotype AI 519 showed specific adaptation to location 1. Except them all are showing better adaptability all three tested environments (Fig.2). Variance for the phenotypic traits indicated that the germplasm selected for the present study were significantly different from each other. The descriptive statistics for the kernel and cob parameters implied the prevalence of good amount of variation in the genotypes (Table.3), which were targeted for the final selection of KRN lines. The kernel row number ranged from 10 to 26 and cob length varied from 5.00 cm to 29.67cm. Another parameter relating to and valid to the present concept is cob girth. It is one of the important traits exhibiting a positive correlation with the KRN and values ranged from 13.08 mm to 58.00 mm. Similarly, kernel per row (8.00 to 41.00) and test weight (6.40 to 40.40 g) also showed high degree of variability. Grouping based on phenotypic diversity formed seven clusters based on 40% dissimilarity index (Fig. 1). In cluster II maximum inbred lines (13) were present followed by clusters III and VI both having 11 each inbred line. While, clusters I and IV had four inbred lines each, clusters V and VII comprised of solitary genotype.

**Fig:1.**
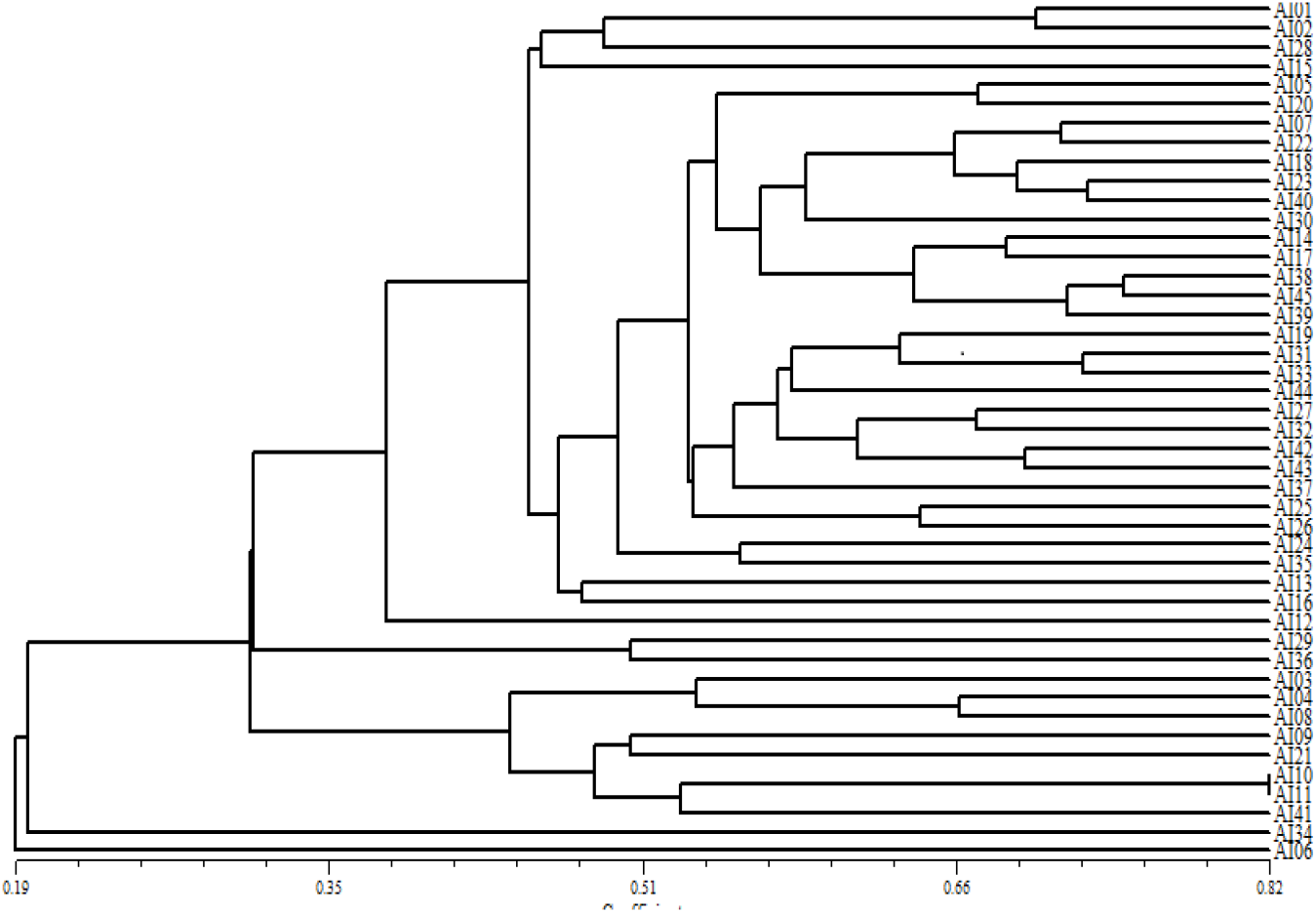
Clustering of genotype based on the morphological diversity

**Fig.2 :**
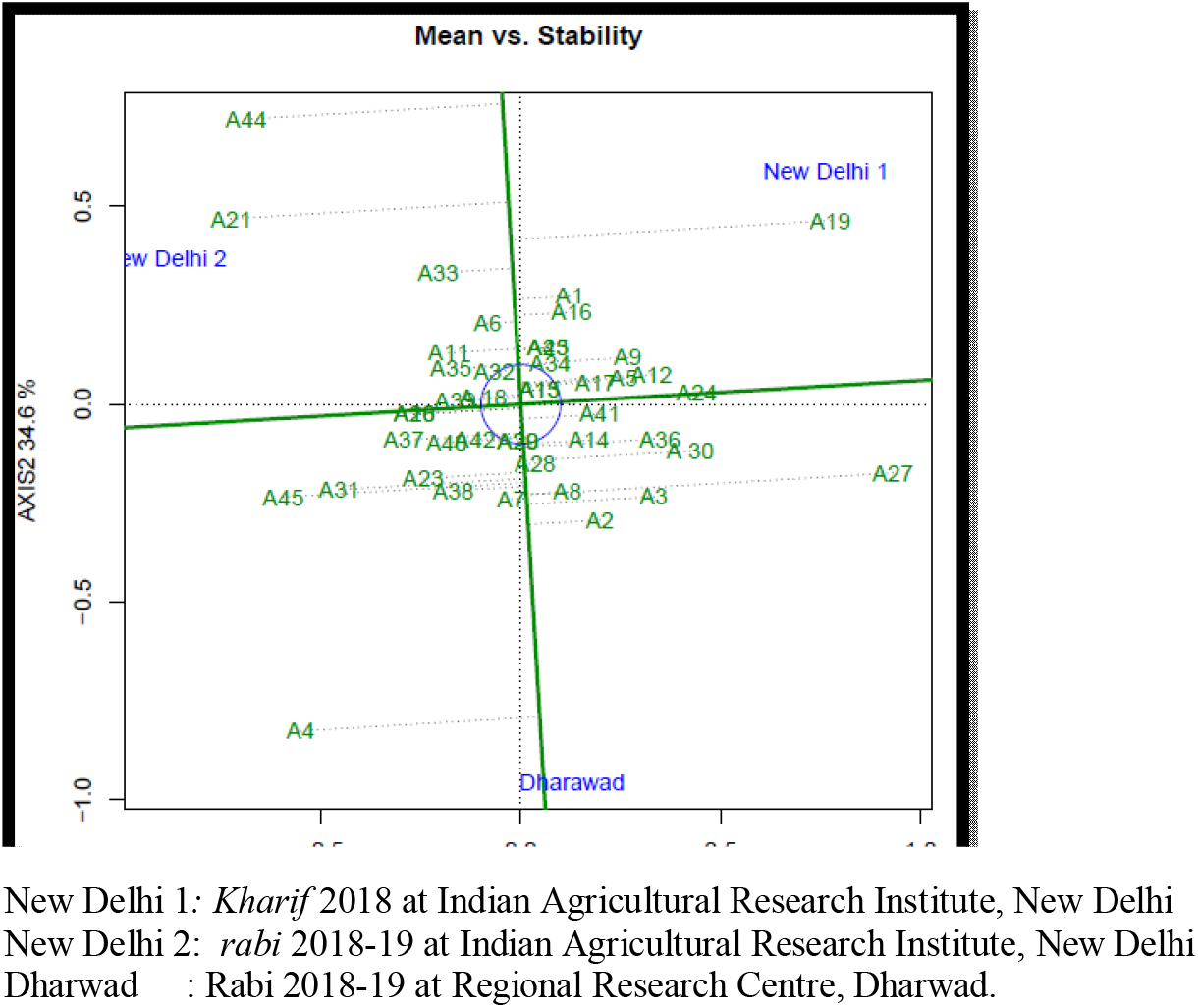
Average environment coordination (AEC) view of GGE bi-plot based on environment focused scaling for the mean performance vs. adaptability for kernel row number across three environments

**Table: 2.**
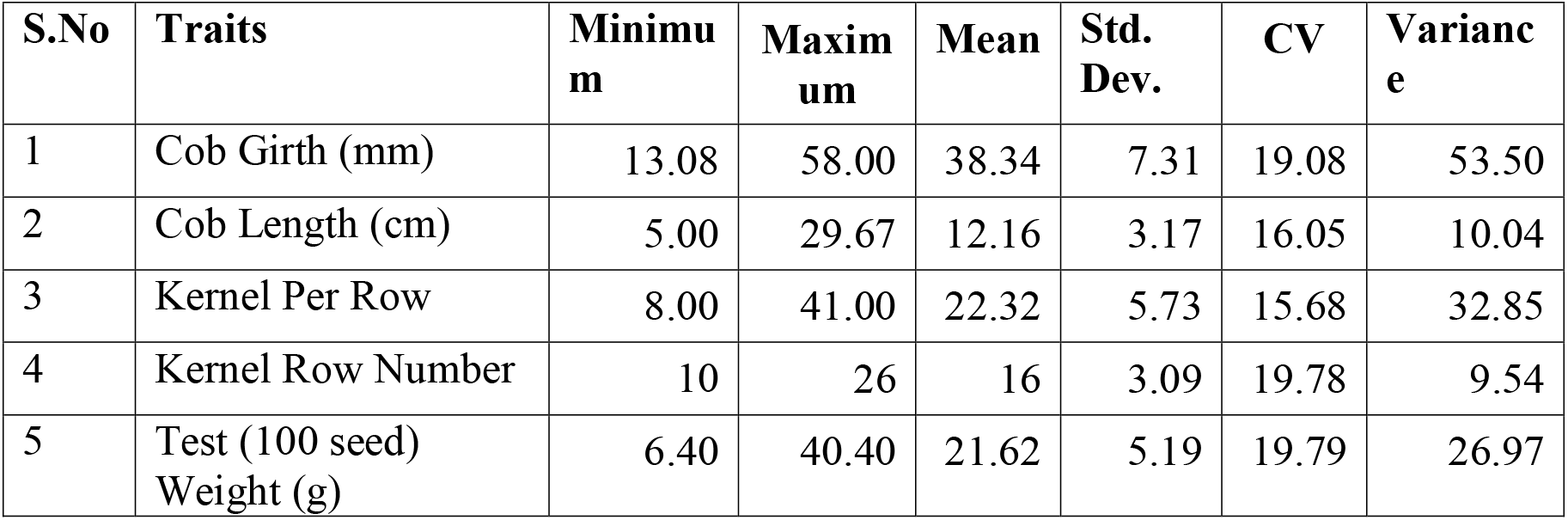
Variability for yield component traits selected 45 tropical maize inbred lines

| S.No | Traits | Minimum | Maximum | Mean | Std. Dev. | CV | Variance |
| --- | --- | --- | --- | --- | --- | --- | --- |
| 1 | Cob Girth (mm) | 13.08 | 58.00 | 38.34 | 7.31 | 19.08 | 53.50 |
| 2 | Cob Length (cm) | 5.00 | 29.67 | 12.16 | 3.17 | 16.05 | 10.04 |
| 3 | Kernel Per Row | 8.00 | 41.00 | 22.32 | 5.73 | 15.68 | 32.85 |
| 4 | Kernel Row Number | 10 | 26 | 16 | 3.09 | 19.78 | 9.54 |
| 5 | Test (100 seed) Weight (g) | 6.40 | 40.40 | 21.62 | 5.19 | 19.79 | 26.97 |

**Table 3.** Pooled ANOVA for cob and kernel parameters across three environments

| Source | Df | Cob length | Cob girth | Kernel row number | Kernel/row | Grain yield |
| --- | --- | --- | --- | --- | --- | --- |
| Genotypes | 44 | 375.5** | 2.3** | 59.68** | 3.0287** | 1.536** |
| Environments | 2 | 555.2** | 0.3691 | 10.12** | 0.607** | 5.489** |
| Interaction | 88 | 137.1** | 0.2091** | 2.91** | 0.1729 | 0.378** |
| Error | 132 | 12.2 | 0.0765 | 0.33 | 0.0963 | 0.025 |

**Table:4.**
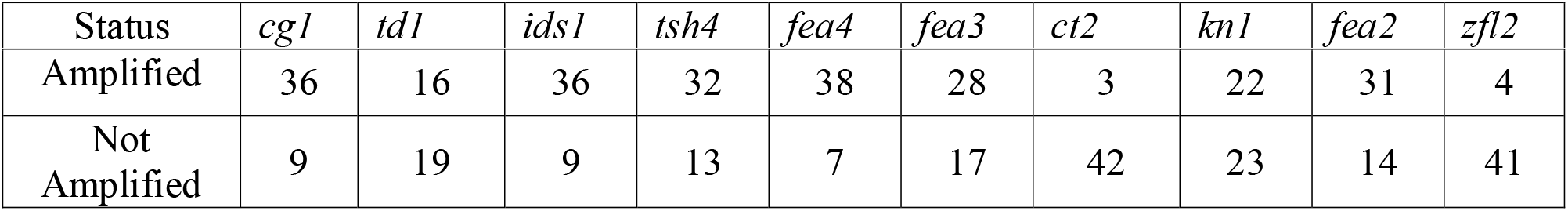
Amplification status of KRN gene across tested maize genotypes

| Status | *cg1* | *td1* | *ids1* | *tsh4* | *fea4* | *fea3* | *ct2* | *kn1* | *fea2* | *zfl2* |
| --- | --- | --- | --- | --- | --- | --- | --- | --- | --- | --- |
| Amplified | 36 | 16 | 36 | 32 | 38 | 28 | 3 | 22 | 31 | 4 |
| Not Amplified | 9 | 19 | 9 | 13 | 7 | 17 | 42 | 23 | 14 | 41 |

### Allelic variability for KRN

Of the 13 genes tested, 10 got amplified in the selected set of maize germplasm subjected to finer characterization. Pattern of amplification profiles of genes as an example in the maize inbred lines is depicted in Fig.3. Observation of amplification indicated that, out of 45 genotypes, 38 showed amplification of *fea4* gene, which is maximum compared to other genes while *cg1* and *ids1* got amplified in 36 genotypes, the remaining nine genotypes had no amplification. Of the other genes *td1, tsh4, fea3, ct2, kn1, fea2* and *zfl2* got amplified in 16, 32, 28, 3, 22, 21, and 04 genotypes, respectively. Interestingly the genotype in which amplification was successful, have also been showing variability for KRN and they were categorized as high (≥14 KRN) and low KRN genotypes. Accordingly, *cg1* is present in 31 high and five low KRN genotypes. The gene *td1, ids1, tsh4, fea4, fea3, ct2, kn1, fea2, zfl2* were present in 14, 30, 26, 30, 25, 3, 17, 25 and 03 high KRN genotypes respectively and also, they were present in 02, 06, 06, 08, 03, 0, 05, 06 and 01 low KRN genotypes respectively (Fig.3). On the other hand, observation of amplification in the genotypes under investigation also had null to multiple genes with them (Table 5). Of the 45 genotypes, only AI 514 showed no amplification whereas nine genes were amplified in two genotypes, *viz*., AI 506 and AI 538. Eight genes got amplified in AI 522, PML 46, AI 517, AI 520, AI 531, AI 533 and AI 539. Similarly, seven genes were amplified in the genotypes, AI 523, AI 526, AI 536, AI 537, AI 519 and AI 527. With six genes were amplification in AI 502, AI 518, AI 521, AI 534, PML 102, AI 503, AI 524, AI 525, AI 528 and AI 529 this group was biggest comprising 10 genotypes. In seven genotypes viz., AI 511, AI 530, AI 535, AI 507, AI 513, AI 532 and PML 17 total of five genes got amplified, whereas four genotypes (*viz*., AI 516, AI 504, AI 505 and AI 509) showed amplification of four genes. While AI 501, PML 95 showed amplification of three genes, in three genotypes (*viz*., AI 510, AI 540 and AI 508) exhibited amplification of two genes responsible for KRN in maize. Only single gene amplification was observed in the genotypes, AI 512 and PML 93.

**Fig:3.**
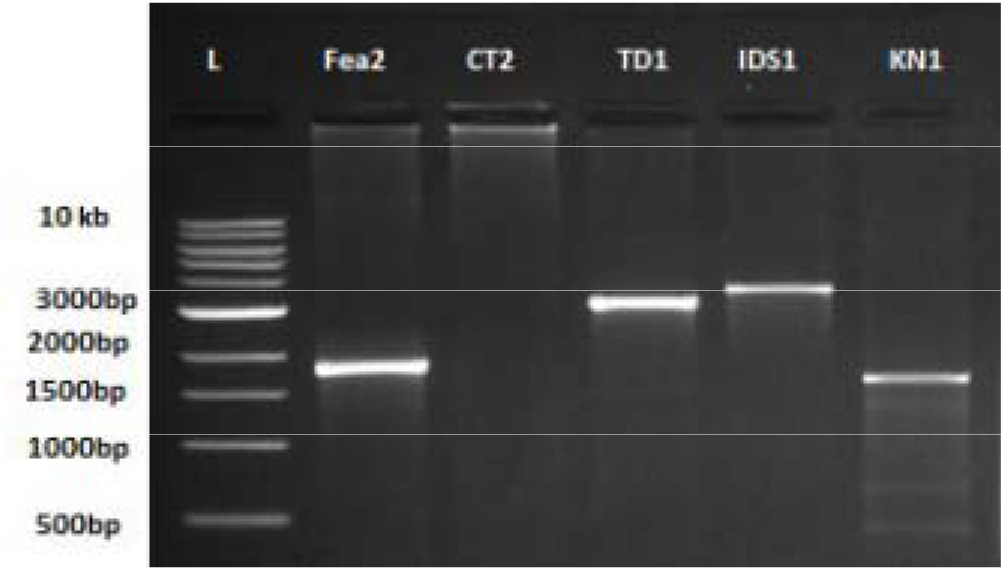
Gel image depicting the amplification of the genes in tropical maize inbred lies

**Table: 5.** Genotypes carrying multiple genes responsible for KRN

| Number of genes |  |  |  |  |  |  |  |  |  |
| --- | --- | --- | --- | --- | --- | --- | --- | --- | --- |
| Null | 1 | 2 | 3 | 4 | 5 | 6 | 7 | 8 | 9 |
| AI 514 | AI 512,PML 93 | AI 510,AI 540,AI 508 | AI 501,PML 95, | AI 516,AI 504,AI 505,AI 509 | AI 511,AI 530,AI 535,AI 507,AI 513,AI 532,PML 17 | AI 502,AI 518,AI 521,AI 534,PML 102,AI 503,AI 524,AI 525,AI 528,AI 529 | AI 523,AI 526,AI 536,AI 537,AI 519,AI 527, | AI 522,PML 46,AI 517,AI 520,AI 531,AI 533,AI 539, | AI 506,AI 538 |

### Influence of Kernel row number by KRN related gene

The amplification of the gene in a genotype was scored as “+” or based on their presence or absence respectively (Supplementary Table 3). These observations were converted in to binary variables and subjected them to logistic regression analysis to explore and understand the probability of KRN gene that influences the kernel row number in these maize genotypes. The coefficients of logit regression were computed and observed values for the gene *cg1, td1, ids1, tsh4, fea3, ct2, kn1, fea2* and *zfl2* were 2.56, 2.67, −0.93, −082, 2.08, −1.32, −1.79, 1.79 and −1.66 with the standard error 1.43, 1.24, 1.42, 1.22, 1.21, 1.61, 0.98,1.13 and 1.13 respectively. Among them gene, *cg1, td1, fea3, kn1* and *fea2* were found significant. Similarly, marginal effects on the gene *cg1, td1, ids1, tsh4, fea3, ct2, kn1, fea2* and *zfl2* were 0.47, 0.58, −0.23, −0.20, −0.48, −0.28, −0.42, 0.39 and −0.34 with the standard error 0.16, 0.20, 0.33, 0.29, 0.23, 0.27, 0.20, 0.19 and 0.23 respectively with Log likelihood −20.25. Thus, comparatively the following genes *cg1, td1, fea3, kn1* and *fea2* had significant marginal effect (Table 6).

**Table:6.**
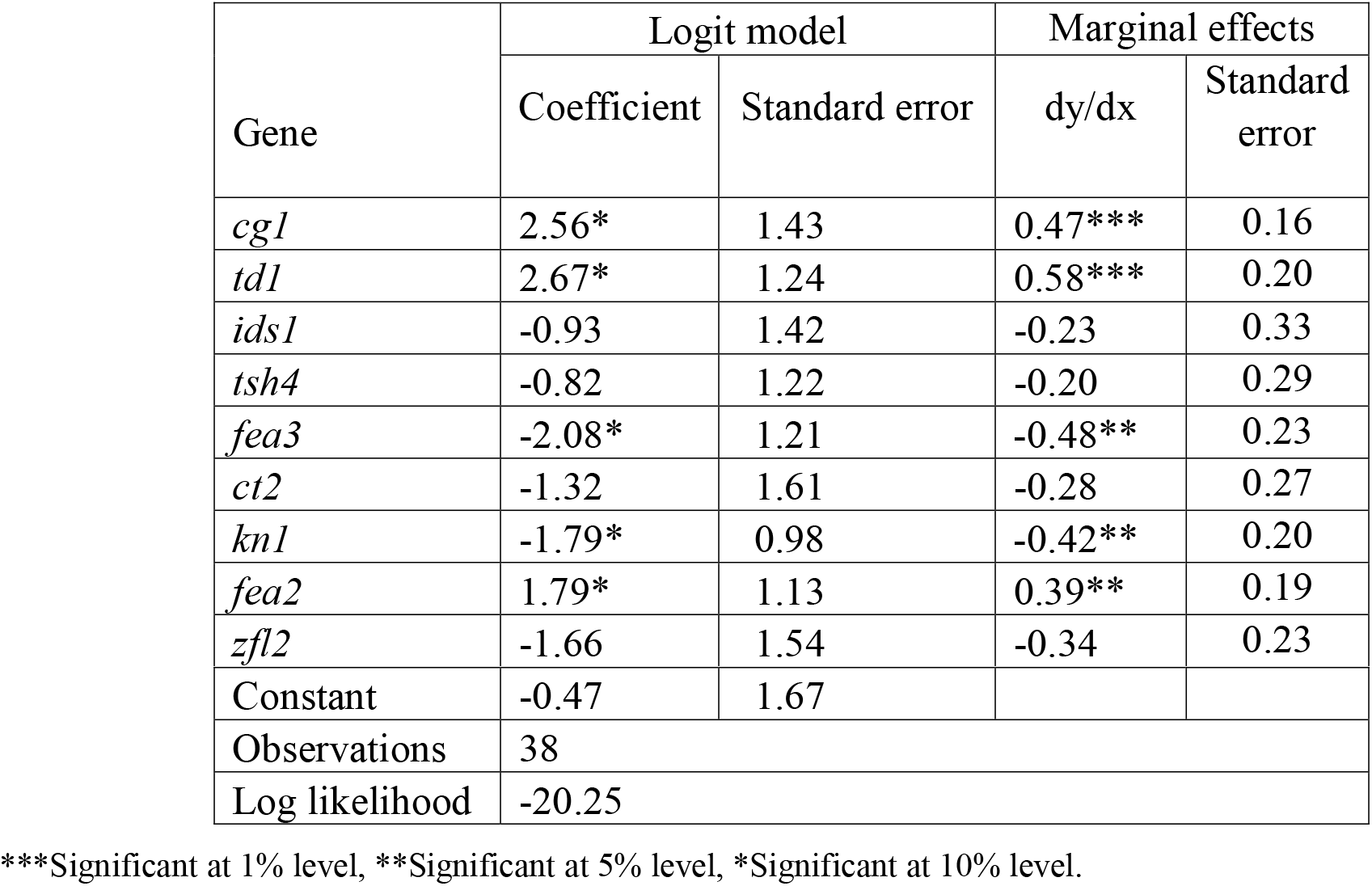
Maximum likely-hood of KRN genes influencing KRN trait in tropical maize inbred line

## Discussion

Maize kernel row number (KRN) per ear is one of the most important yield components and is a breeding goal for the improvement of maize inbred lines. To better understand the grain yield, QTLome, dozens of QTLs and trait-associated variants (TAVs) for kernel row number (KRN) have been identified across the maize genome by linkage mapping and next-generation sequencing-based genome-wide association studies (Li et al. 2018; Xiao et al. 2017; Calderón et al. 2016; Liu et al. 2015a; Peng et al. 2011; Lu et al. 2010; Messmer et al. 2009; Upadyayula et al. 2006; Veldboom and Lee 1994; Stuber et al. 1987). Further the knowledge of genetic architecture of KRN was utilized to establish breeding programs. But the systematic study in unraveling the genetic base of the yield components targeting kernel row number is lacking in tropical maize. In the present study an effort was made to characterize and select a set of tropical maize inbred lines from the pool of germplasm and further understand the association of KRN related genes to the kernel row number. The selected 45 inbred lines for the current study was significantly different from each other as per kernel row per ear were concerned and adaptable to varied environments tested. The expression of kernel row number was not much affected by the environment in which genotypes were grown. The range of KRN (10 to 26) in the studied maize inbred lines indicated its potential variability harnessed in crop improvement program. Along with the KRN, it was observed that variance parameters for cob girth, cob length, kernel per row, test weight were also high, indicating the diverse nature of the inbred lines. Diversity analysis among the selected 45 inbred lines also showed possible cause for this variation and grouping of the genotypes were done majorly based on the KRN and related component traits. Taking a clue from these two analyses, inbred lines were characterized for the genes responsible for KRN. The length of the gene under characterization was ranged from 1573 bp to 7928 bp with varied exon and intron numbers. Amplification profile of 13 genes indicated (Supplementary Table 3) that majority of the genes are present in Indian maize inbred line also, a single copy of gene amplified as per the expected amplicon size. Only three genes which could not amplify in the studied material may not be present in the inbred lines or they may be specific to the genetic background of the maize inbred lines (Chuck et al., 2014). The genotypes which carried gene of interest were classified in to high and low KRN types. It indicated that, the studied genes were having high a tendency to move with high KRN genotypes compared to their low KRN counterparts (Fig.4). This implies that majority of the genes taken in this study are associated with high KRN phenotype (Bommert et.al, 2005 and 2013, Chucket.al, 2007 and 2010, Bomblies et.al, 2006, McSteen et.al. 2006). The genetic control of kernel row number showed estimates that allow the researcher to infer that this is a polygenic character with predominantly additive allelic interaction (Toledo et al., 2011). It is evident from the study that Indian maize germplasm also contains one or more KRN related genes for its phenotypic expression as high KRN genotype (Table 5). Recently, several genes in the CLV-WUS feedback loop, including thick tassel dwarf1 (td1) (Bommert et al, 2005), fasciated ear2 (fea2) (Taguchi-Shiobara et al. 2001), and COMPACT PLANT2 (CT2) (Bommert et al. 2013) were isolated in maize. Additionally, the RAMOSA genes (McSteen et al. 2006) Corngrass1 (Cg1) (Chuck et al. 2007), tasselsheath4 (tsh4) (Chuck et al, 2010), FLORICAULA/LEAFY (ZFL1 and ZFL2) (Bomblies et al. 2006), unbranched2 (ub2) and ub3 (Chuck et al. 2014) and others, all affect ear morphology by regulating the development of SPMs and SMs which finally decides the KRN in maize. The marginal effects of maximum likelihood indicated that *cg1, td1* and *fea2* are promoting high kernel row number by 47%, 58% and 39% respectively. It may be inferred that if *cg1* is present in the maize inbred lines, the probability of getting a high kernel row number is 47 %. Similar examinations hold good for the other two genes also. Natural variation in *fea2* explains part of the difference in KRN between inbreds Mo17 and B73 (Bommert et al. 2013). The *fea2* and *td1* are proposed to regulate *ZmWUS1* based on the well-known model in Arabidopsis.

**Fig: 4.**
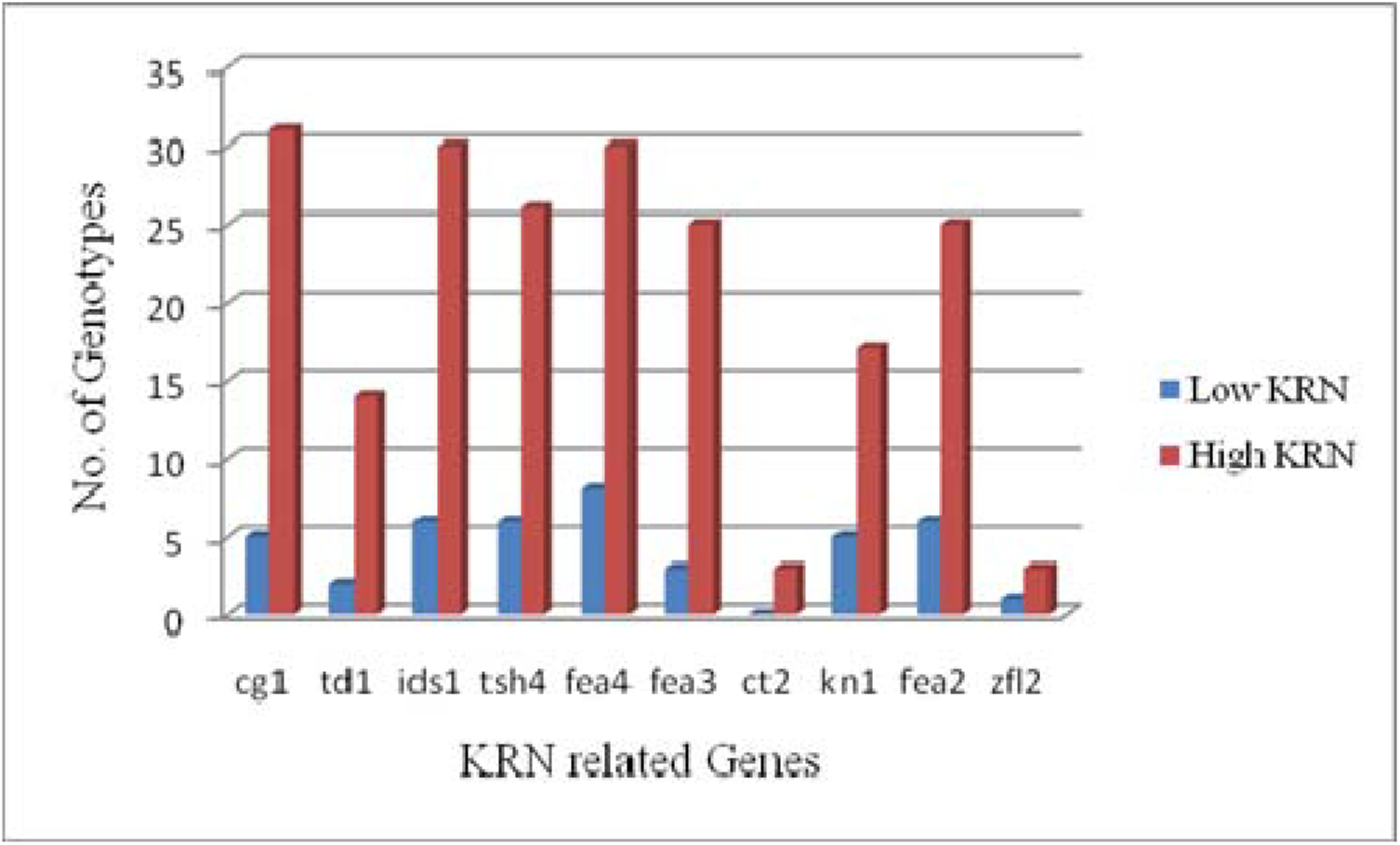
Tendency of KRN genes co-developing with high and low KRN genotypes

**Fig. 5.**
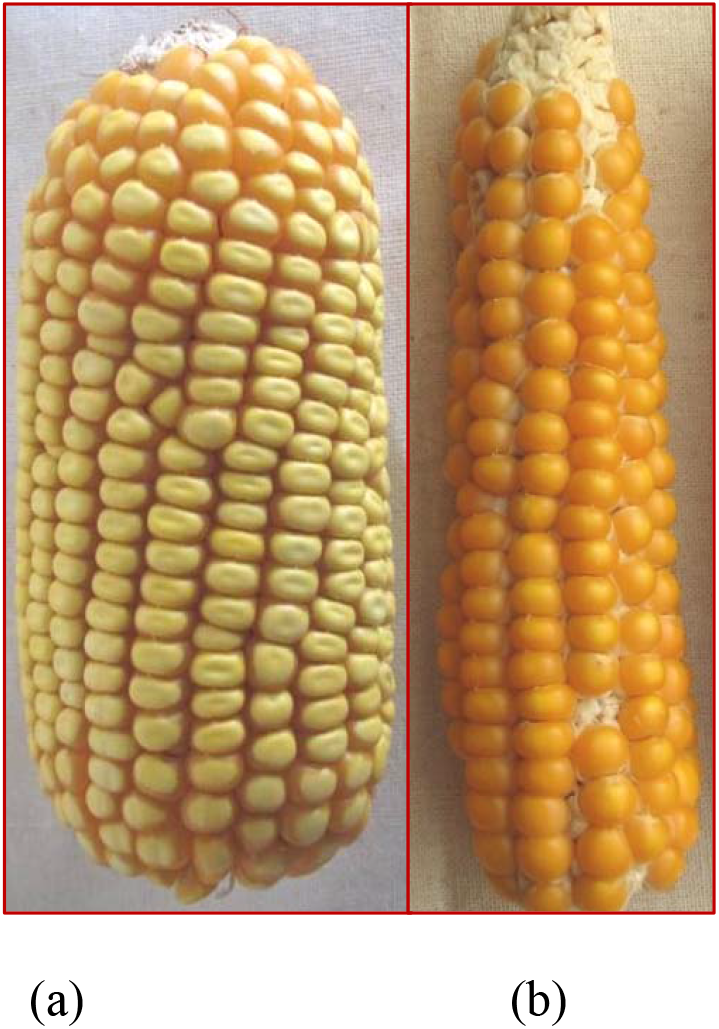
Phenotype of high (a) and low (b) kernel genotype

On the other hand, if *fea3* gene is present in the maize inbred lines, there will be a 48 % probability that *fea*3 will reduce the KRN in maize. It influences the addition of kernel row number in a negative direction. The earlier experimentation with *Fasciated* mutant ears, such as those produced by *fea3* mutants, have low seed yield, because they make too many disorganized primordia that do not fill properly. However, it was found that weak alleles of *fea3* make more kernels (Byoung et al, 2016). Hence allelic variation in the *fea3* may be dissected to understand the role of *fea3* gene in tropical maize inbred lines. Further, genes with monomorphic amplification among the contrast genotypes may be studied in details to understand its association with kernel row number in maize.

## Conclusion

Enhancing kernel row number in maize is one of the important breeding targets in maize in general and for hybrid developing strategy in particular. The KRN genes which are characterized in temperate germplasm may have background effect and possibly available in different allelic forms tropical maize inbred lines. However, four genes, *cg1, td,* fea2 and *fea3* showed association with KRN in the tropical maize inbred lines. The findings can be further studied and confirmed through molecular expression analysis.

## Acknowledgements

This study is the output of an external project funded by DST-SERB, New Delhi. Authors are thankful for the financial support by the DST, New Delhi

## Compliance with Ethical Standards

### Funding

This study was funded by DST-SERB, New Delhi, India with the grant No.EEQ/2016/000447. This grant was received by author^1^.

### Ethical approval

This article does not contain any studies with human participants or animals performed by any of the authors. Hence it doesn’t require any ethical approval.

---

**Supplementary Table 2.:**
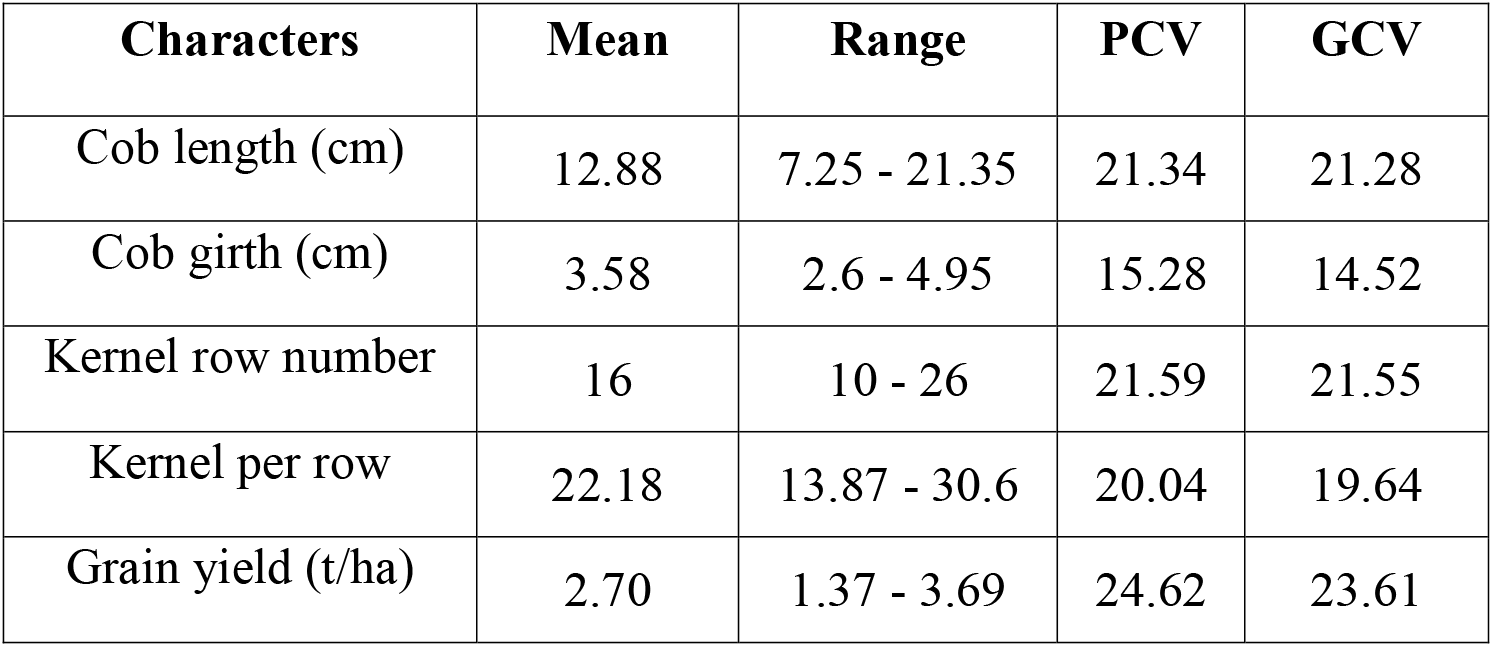
Genetic variability parameters for selected 80 maize inbred lines

**Supplementary Table 3.**
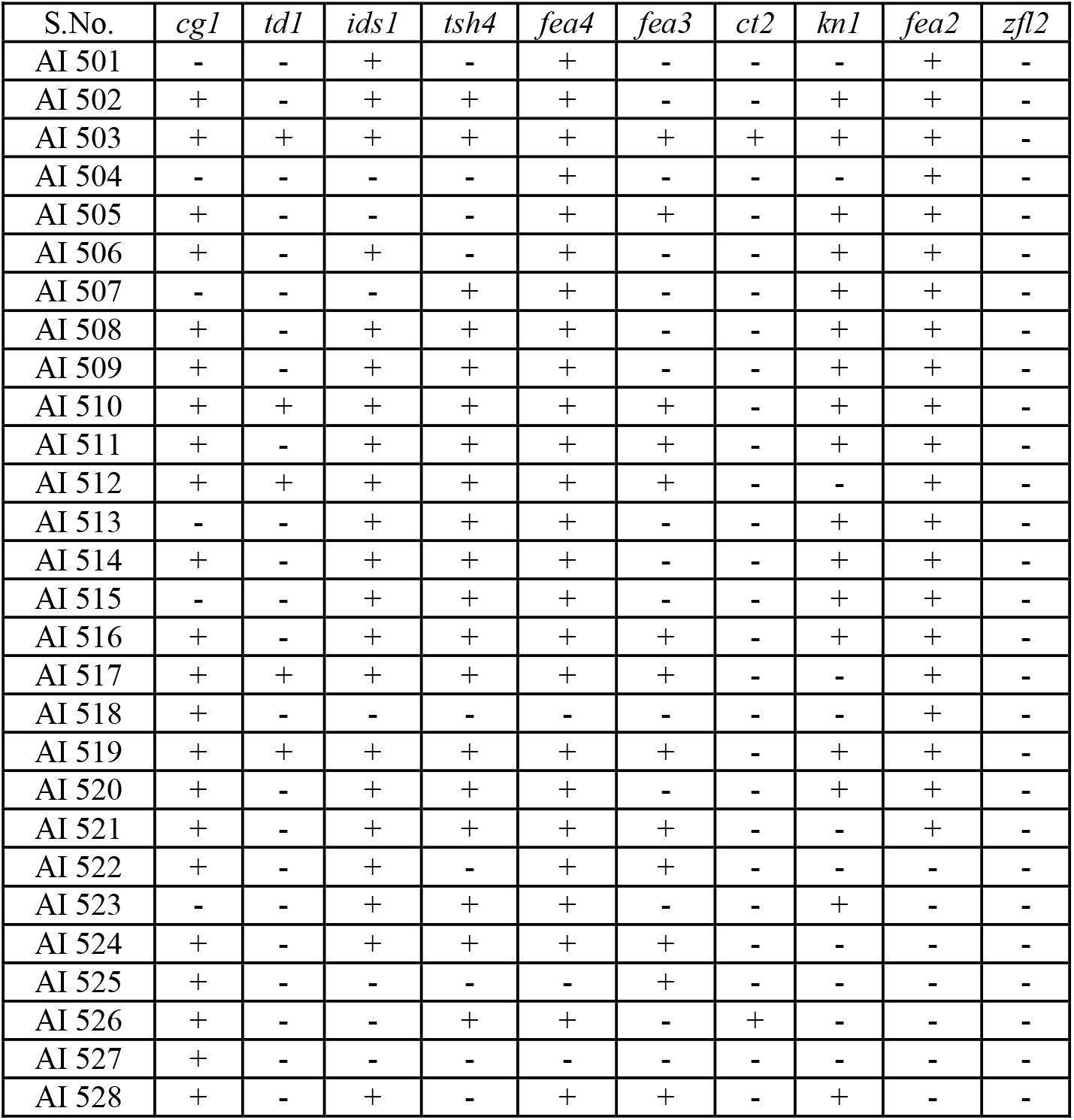
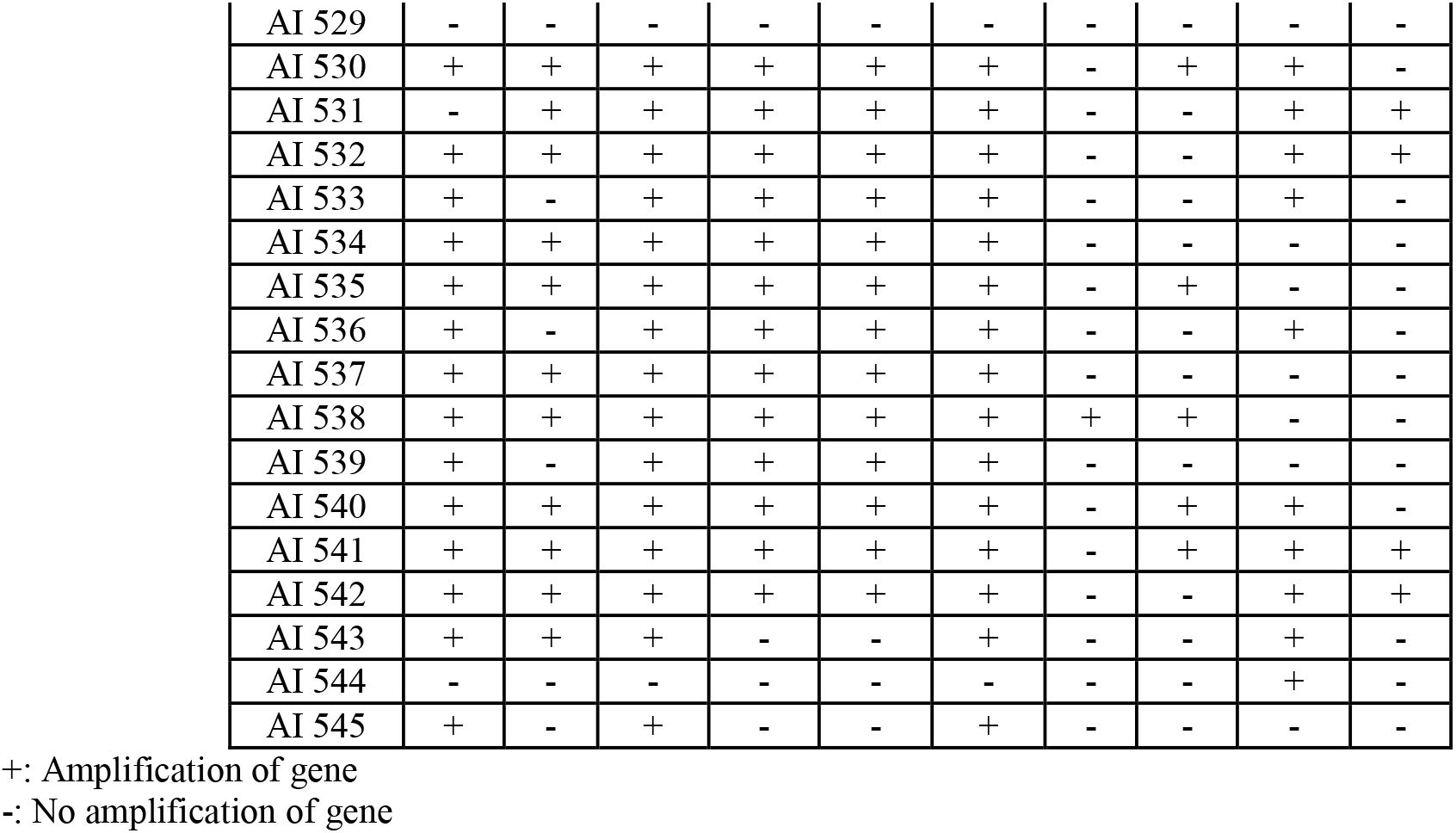
Gene amplification profile of KRN related genes in tropical maize inbred lines

**Supplementary Table 1:** Mean and range for cob and kernel traits in 280 tropical maize germplasm

| Variables/<br>Season | Cob Length (cm) |  | Cob Girth (mm) |  | Kernel Row<br>Number |  | Kernels/ Row |  | Shelling Percentage |  |
| --- | --- | --- | --- | --- | --- | --- | --- | --- | --- | --- |
|  | Kharif<br>2017 | Rabi 2017-18 | Kharif<br>2017 | Rabi<br>2017-18 | Kharif<br>2017 | Rabi<br>2017-18 | Kharif<br>2017 | Rabi<br>2017-18 | Kharif 2017 | Rabi<br>2017-18 |
| Mean | 13.21 | 12.16 | 3.12 | 3.22 | 14 | 14 | 26.68 | 23.18 | 67.65 | 69 |
| Range | 3.2-20.80 | 6.75-19.60 | 1.50-4.40 | 2.05 - 4.75 | 10-26 | 10-26 | 21-46 | 14-36 | 52-90 | 56-90 |
| Std Dev | 2.65 | 2.90 | 0.53 | 0.60 | 2.42 | 2.50 | 5.70 | 5.60 | 10.19 | 11.00 |

